# Genetic modification of the bee parasite *Crithidia bombi* for improved visualization and protein localization

**DOI:** 10.1101/2024.01.18.576273

**Authors:** Blyssalyn V. Bieber, Sarah G. Lockett, Sonja K. Glasser, Faith A. St. Clair, Lynn S. Adler, Megan L. Povelones

## Abstract

*Crithidia bombi* is a trypanosomatid parasite that infects several species of bumble bees (*Bombus* spp.), by adhering to their intestinal tract. *Crithidia bombi* infection impairs learning and reduces survival of workers and overwintering queens. Although there is extensive research on the ecology of this host-pathogen system, we understand far less about the mechanisms that mediate internal infection dynamics. *Crithidia bombi* infects hosts by attaching to the hindgut via the flagellum, and one previous study found that a nectar secondary compound removed the flagellum, preventing attachment. However, approaches that allow more detailed observation of parasite attachment and growth would allow us to better understand factors mediating this host-pathogen relationship. We established techniques for genetic manipulation and visualization of cultured *C. bombi*. Using constructs established for *Crithidia fasciculata*, we successfully generated *C. bombi* cells expressing ectopic fluorescent transgenes using two different selectable markers. To our knowledge, this is the first genetic modification of this species. We also introduced constructs that label the mitochondrion and nucleus of the parasite, showing that subcellular targeting signals can function across parasite species to highlight specific organelles. Finally, we visualized fluorescently tagged parasites *in vitro* in both their swimming and attached forms, and *in vivo* in bumble bee (*Bombus impatiens*) hosts. Expanding our cell and molecular toolkit for *C. bombi* will help us better understand how factors such as host diet, immune system, and physiology mediate outcomes of infection by these common parasites.

## Introduction

Trypanosomatids of the class Kinetoplastea are single-celled eukaryotic parasites [1]. While some trypanosomatid species, such as *Leishmania*, can be transmitted to humans by insect vectors causing considerable morbidity and mortality [2], most trypanosomatids are monoxenous and exclusively parasitize insects [3]. For these parasites, fecal-oral transmission is the most common mode of pathogen spread [4]. This cycle requires an infected host to defecate parasites onto a food source, where they must remain viable long enough to be ingested by their next susceptible host. Once in the host, parasites often accumulate in the hindgut and rectum, adhering to the lining of these tissues by their single flagellum and dividing by binary fission as attached cells [4]. The structure of this flagellar attachment is similar in all trypanosomatids [5,6].

Some trypanosomatids infect bees such as honey bees (*Apis mellifera*) and bumble bees (*Bombus* spp.), potentially contributing to pollinator decline [7]. For example, *Crithidia bombi* is a gut parasite primarily known to infect multiple species of bumble bees, including *Bombus impatiens* and *Bombus terrestris*, although *C. bombi* has recently been found to replicate in the solitary bee species *Osmia lignaria* and *Megachile rotundata* as well [8]. In bumble bees, *C. bombi* impairs learning [9], can reduce queen colony-founding success [10], and can reduce worker survival under stressful conditions [11]. Other trypanosomatids, such as *Lotmaria passim*, cause similar effects in honey bees [12]. The presence of the parasites triggers an innate immune response in the bee host, although precisely how infection impacts host fitness is unclear [13,14]. Researchers have also shown that gene expression patterns in cultured parasites differ from those of parasites in the bee gut, representing possible metabolic adaptations to the host environment [14].

Certain floral diets can reduce *C. bombi* infections in some bumble bee species. The secondary metabolite callunene, discovered in the nectar of heather flowers (*Calluna vulgaris*), removed or shortened the flagellum of *C. bombi* and dramatically reduced infection in *B. terrestris*, presumably by interfering with the parasites’ ability to adhere to and colonize the gut [15]. Similarly, pollen of sunflower (*Helianthus annuus*) and some other Asteraceae plants dramatically decreases *C. bombi* infection in *B. impatiens* [16–20] but is less effective in other *Bombus* species [21], suggesting that species-level variation shapes diet-mediated effects on infection outcomes.

The underlying mechanisms for the antiparasitic effect of different floral products such as pollen and nectar are largely unknown. Molecular genetic tools to manipulate parasites for *in vivo* infections and in culture would facilitate new experimental approaches to understand how floral resources impact host-pathogen dynamics. For instance, which parasite biological processes are disrupted by heather nectar or sunflower pollen? Possible targets include flagellar growth, attachment, and survival and division of attached cells. Discovering the effects of floral products on these activities could improve our understanding of how these different aspects of parasite biology contribute to productive infections. Although all trypanosomatids, including human pathogens, attach to tissues in their insect hosts [6], insect parasites do so in great numbers [4], meaning they could serve as a model for insect colonization by trypanosomatids more generally. In addition, improved understanding of the effects of pollen and nectar diets on the mechanisms underlying parasite infections could allow us to predict the impacts of floral resources on pathogen load and pollinator health.

Detailed study of attachment and modes of cell division would be greatly facilitated by improved visualization of parasites *in vivo*. For this, both whole cell and organelle markers would allow researchers to monitor the number, location, and cellular structure of parasites at different stages of the infection. Such analyses would improve our understanding of the life cycle of these parasites in their insect hosts, which could reveal vulnerabilities for intervention. *In vitro* assays for infection behaviors such as attachment would allow for time-resolved, quantitative studies showing how attachment changes under different conditions, predicting infection dynamics in the presence and absence of different floral products and compounds. Finally, genome-wide transcriptomics and proteomics approaches will reveal gene products that mediate interactions between parasites and their insect hosts [13]. Genetic techniques enabling functional knockout and subcellular localization during the cell and life cycle of the parasite would provide important insights into the mechanism of action of specific proteins.

To develop these approaches, our objectives for this study were to 1) establish *C. bombi* sensitivity to antibiotics used as selectable markers, 2) introduce episomal plasmids including genes for enhanced green fluorescent protein (eGFP) and red fluorescent protein (RFP) into *C. bombi* cells, 3) create markers for subcellular organelles, 4) isolate and culture parasites from *B. impatiens* intestinal tracts, and 5) visualize fluorescently-labelled *C. bombi* cells *in vitro* and *in vivo*.

## Methods

### Parasite lines and culture

To establish sensitivity to antibiotics, *C. bombi* strains 08.076 and 16.075 (provided by Ben Sadd, Illinois State University) were cultured in FP-FB media supplemented with 2 μg/mL hemin (Sigma, St. Louis, MO) and 10% fetal bovine serum (Atlanta Biologicals, Bio-Techne, Minneapolis, MN) at 27 °C and 3% CO_2_, as described [22]. For drug sensitivity tests and maintaining genetically modified parasite lines, medium was supplemented with either hygromycin (Hyg, catalog number 10687010, ThermoFisher) or neomycin (Neo, G418, catalog number G8168, Sigma). Growth curves were performed in triplicate and drug concentrations ranging from 0 to 80 μg/ml in 2-fold increments were tested. Parasites were grown in sterile, untreated tissue culture plates or 25 cm^3^ flasks with vented caps. Cell densities were determined by removing a 25 μL sample of the culture to a 1.5 mL tube, adding 25 μL of 3% formalin to fix, followed by 200 μL of Gentian violet (Harleco) staining solution [23]. 10 μL of this mixture was then applied to a Neubauer hemacytometer and counted on an inverted tissue culture light microscope (Zeiss Primovert). Parasites were maintained between 5 x 10^5^ and 5 x 10^7^ cells/mL by diluting in fresh medium every 2-3 days. To generate attached parasites, 2 mL of log-phase parasites (1-2 x 10^7^ cells/mL) were allowed to adhere to a poly-L-lysine coated dish (MatTek, Ashland, MA) for 24 hours, followed by washing 3X with 1X PBS to remove non-attached cells.

### Plasmids and transfection

To introduce plasmids into *C. bombi*, the pNUS series of plasmids containing sequences for expression of transgenes in *Crithidia fasciculata* and *Leishmania* were used (provided by Emmanuel Tetaud) [24]. The pNUS-eGFP-cH (enhanced green fluorescent protein, hygromycin resistance) and pNUS-RFP-cN (red fluorescent protein, neomycin resistance) were transformed unmodified into *C. bombi* strains 08.076, 16.075, or WHA1. For organelle markers, the pNUS-mitoeGFP-cH was created as described [25]. To create pNUS-*Cf*RNH1eGFP-cH, the open reading frame (lacking the stop codon) for the RNase H1 gene from *C. fasciculata* (*Cf*RNH1, TriTrypDB accession number CFAC1_220025400) was amplified by PCR using primers 5’-GCTACTAGCATATGATGAAGCCGTCGTTTTATGTA and 5’-GCTACTAGGGTACCCTCACTGGTCCCGTGCATACG containing NdeI and KpnI restriction enzyme sites, respectively (underlined). The amplified product was cloned into pNUS-eGFP-cH using NdeI and KpnI restriction sites resulting in fusion of eGFP to the C-terminus of *Cf*RNH1. Plasmids were confirmed by Sanger (Eurofins Genomics, Louisville, KY) or Nanopore (Plasmidsaurus) sequencing. Circular plasmids were concentrated by ethanol precipitation and resuspended in sterile water at a concentration of 0.5 μg/μL. For nucleofection, 1 x 10^8^ *C. bombi* cells were harvested by centrifugation (5 min, 800 rcf) and resuspended in 100 μL TbBSF buffer [26]. 5 μg of plasmid was added to the tube and the solution was mixed briefly before being transferred to a Lonza cuvette. Parasites were nucleofected using program X-001 on a Lonza Nucleofector 2b device. Cells were then transferred to media and allowed to recover for 24 hours before addition of selecting drug to a final concentration of 40 μg/mL. An equal number of untransformed cells were resuspended in the same drug concentration as a “mock” control for drug effectiveness. After 10 days of selection, surviving cells were screened by fluorescence microscopy. The concentration of selecting drug was then increased to 80 μg/mL.

### Microscopy of cultured parasites

Expression of fluorescent proteins expressed in the cytoplasm or in organelles was detected by fluorescence microscopy. 1 x 10^7^ cells were harvested by centrifugation (5 min, 800 rcf) and washed once with PBS. Cells were allowed to adhere to poly-L-lysine coated coverslips for 20 min in a humid chamber at room temperature followed by 2 washes in PBS. Cells were fixed with cold 4% paraformaldehyde for 15 min then washed twice with PBS. Cells were permeabilized in 0.1% Triton X-100 for 5 min followed by 2 washes in PBS, stained with 2 μg/mL DAPI, washed in PBS, and mounted in Vectashield (Vector Laboratories, Burlingame, CA). Parasites were imaged on a Leica SP8 confocal microscope using the 100x objective. Z-stack images with 0.2 μm steps were taken for all *in vitro* fluorescent cell microscopy. Final images shown in figures are max projections of the z-stack images. Attached parasite rosettes were imaged live in PBS.

### *Isolation of parasites from* Bombus impatiens

*Crithidia bombi* (origin, Hadley, Massachusetts: 42.363911 N, −72.567747 W) were collected from the wild in 2014 and thereafter maintained as a live infection in commercial *B. impatiens* colonies (Koppert Biological Systems, Howell, Michigan, USA and Biobest USA Inc., Romulus, Michigan USA). We randomly selected five bees, dissected their digestive tracts, and homogenized the tissue in 300 μL of isotonic Ringer’s solution. After three hours, a 150 μL sample was taken from the top of the homogenized solution and centrifuged (5 min, 800 rcf). Pelleted material was resuspended in 5 mL of FP-FB medium and stored at 4 °C. Additionally, feces from five infected bees from the same colony were collected using capillary tubes, transferred directly into 5 mL of FP-FB medium, and stored at 4 °C for up to five days. We modified the protocol established in [22] to isolate *C. bombi* cells from the bee digestive tracts and feces. The samples were centrifuged (5 min, 800 rcf) and resuspended in 1 mL of one of two different modified “Mäser Mix” media. One included 1% antibiotic [Penicillin-Streptomycin (Pen-Strep), Sigma Aldrich] while the other included both 1% Pen-Strep and 1% antifungal, Amphotericin B (250 μg/mL, Sigma Aldrich), both in FP-FB medium. Parasites were incubated at 27 ºC and 3% CO_2_ overnight followed by cloning by limiting dilution in 96 well plates at a calculated density of 0.5 cells/well or 0.1 cells/well. Two weeks later, positive clones were identified as containing parasites without bacterial or fungal contaminants and scaled up for further analysis. These 30 clonal isolates were internally named WHA1-WHE6.

### *Laboratory infections of* Bombus impatiens

To visualize *C. bombi* infection *in vivo*, inoculum was prepared by sampling from an early log phase culture (6.5 x 10^5^-6.5 x 10^6^ cells/mL) of *C. bombi* strain WHA1 expressing either eGFP or RFP. To remove cells from media containing selecting drug, cells were centrifuged (5 min, 800 rcf), washed twice with Ringer’s Solution, and then resuspended in equal parts Ringer’s Solution and 30% sucrose solution to create a final inoculum of 1200 cells/μL and 15% sucrose solution. *Bombus impatiens* workers were sampled from uninfected colonies (Koppert Biological Systems, Howell, Michigan, USA), transferred to individual vials, starved for 3-4 hours, then presented with two 15 μL droplets of inoculum and observed until both drops were consumed. Only bees that consumed both droplets were included in experimental trials. To allow for the infection to progress, 5-6 bees inoculated with either eGFP or RFP-expressing *C. bombi* infections were placed into deli cups and fed *ad libitum* on 30% sucrose and wildflower pollen (CC Pollen Company, Phoenix, Arizona, USA). After allowing infections to progress 7 days post-inoculation, we dissected the bees and imaged the digestive tracts using a Nikon TiE microscope with an A1R confocal system equipped with NIS-Elements 5.3 software. Z-stacks with steps 5 μm apart were taken using the 20x objective and final images shown in figures are max projections of the z-stack images.

## Results

### (1) Crithidia bombi *is sensitive to selecting compounds*

Genetic modification requires selection for cells that have taken up exogenous recombinant DNA molecules. In particular, the expression of fluorescent proteins facilitates detailed morphological analysis of parasites both *in vitro* and *in vivo*. Therefore, we sought to establish a protocol whereby plasmids driving expression of fluorescent proteins could be introduced into *C. bombi*. To do this, we first needed to determine sensitivity to antibiotics for which resistance genes could be used as selectable markers. We monitored growth of *C. bombi* strain 08.076 [22] in 0, 5, 10, 20, 40 or 80 μg/ml of either hygromycin (Hyg) or neomycin (Neo, G418) over the course of four days (Fig 1). We found that *C. bombi* were sensitive to both drugs, with 40 μg/mL being the lowest concentration that completely inhibited growth after 20 hours.

**Fig. 1.**
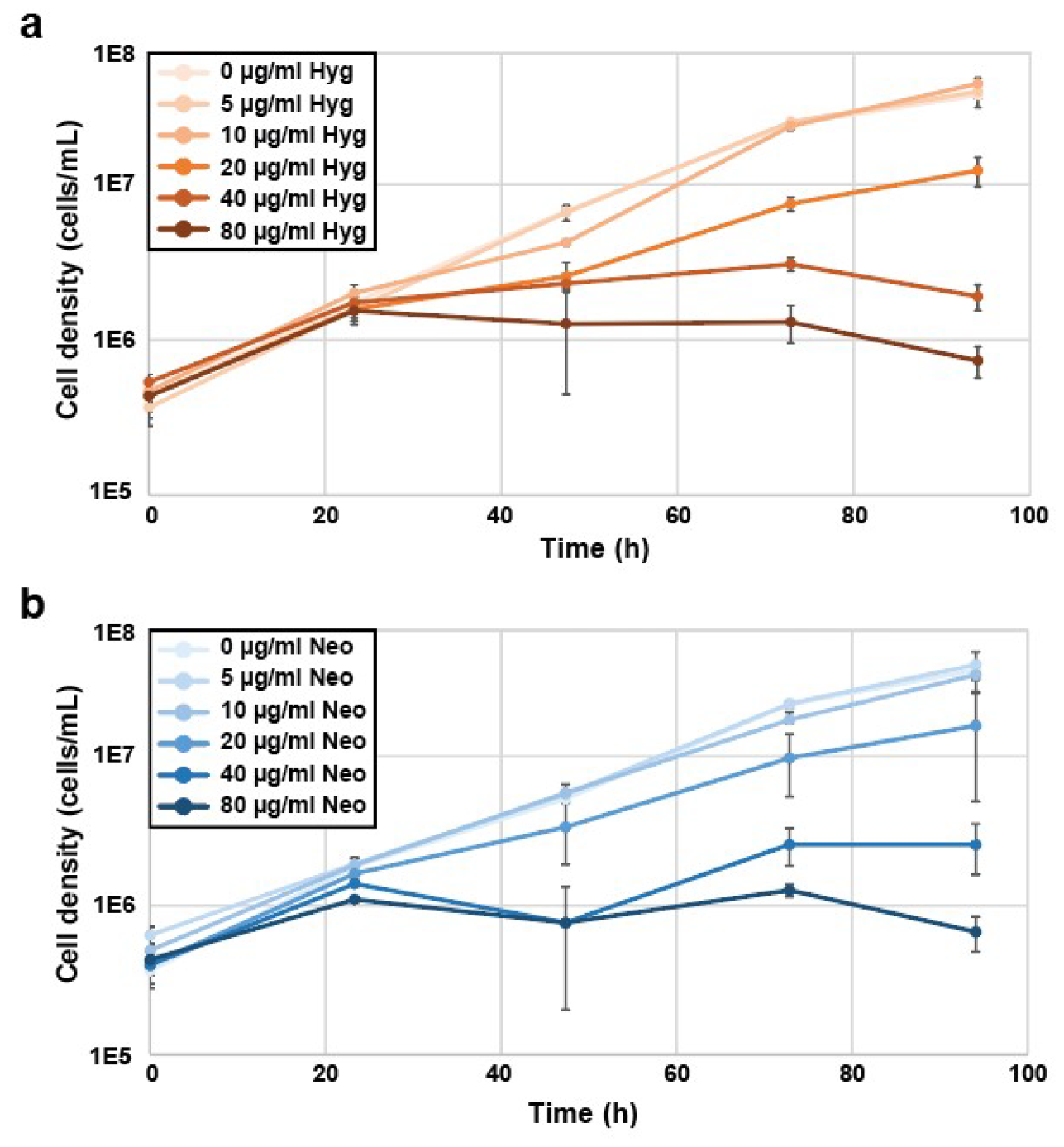
Cultured *Crithidia bombi* cells are sensitive to selecting drugs. Growth curves of *C. bombi* parental strain 08.076 [22] grown in FP-FB medium with increasing levels of **a)** hygromycin (Hyg) or **b)** neomycin (Neo) to determine optimal concentration for selection. Each graph shows the mean of three independent replicates. Error bars are standard error.

### (2) Expression of cytoplasmic fluorescent proteins

Next, we investigated methods for genetic modification of *C. bombi*. Some trypanosomatids can maintain circular plasmids as episomes [27–31]. This approach is advantageous in that it does not require stable integration by homologous recombination at a genomic locus. Using episomes simplifies construct creation, does not require detailed knowledge of the genome sequence, and can increase transformation efficiency. For this reason, we chose a series of plasmids originally developed for the congener *C. fasciculata*, a monoxenous parasite of mosquitoes, and *Leishmania*, a dixenous parasite of sand flies and mammals [24]. The pNUS-eGFP-cH plasmid (Fig 2a) contains genes for eGFP and Hyg resistance flanked by 5’ and 3’ untranslated regions (UTRs) derived from *C. fasciculata* [24]. UTRs direct processing of the mature transcript required for expression [32]. Similarly, pNUS-RFP-cN contains a gene encoding RFP and the Neo resistance gene (Fig 2a). We prepared each of these plasmids by ethanol precipitation and introduced them into the cultured *C. bombi* strain 08.076 by nucleofection using a protocol previously established for *C. fasciculata* [25]. After nucleofection, cells were returned to media in either 24-well plates or in tissue culture flasks and were left to recover for approximately 24 hours before addition of 40 μg/mL of either Hyg or Neo. For each nucleofection, a sample containing the same number of untransformed cells was placed under selection to confirm drug efficacy. After 10-12 days of selection, plates containing nucleofected cells had healthy, dividing cells, while no wells in the mock plates contained growing cells. Since every well in these plates was drug resistant, we conclude that the lines obtained are not clonal. The same result was obtained when cells were selected in flasks. Screening of nucleofected cells by fluorescence microscopy revealed eGFP or RFP expression, indicating successful transformation of the episomal pNUS-eGFP-cH plasmid (Fig 2b) and pNUS-RFP-cN plasmid (Fig 2c). We did not detect any instances of spontaneous Hyg or Neo resistance, as no cells were recovered from the “mock” treatment vessels.

**Fig. 2.**
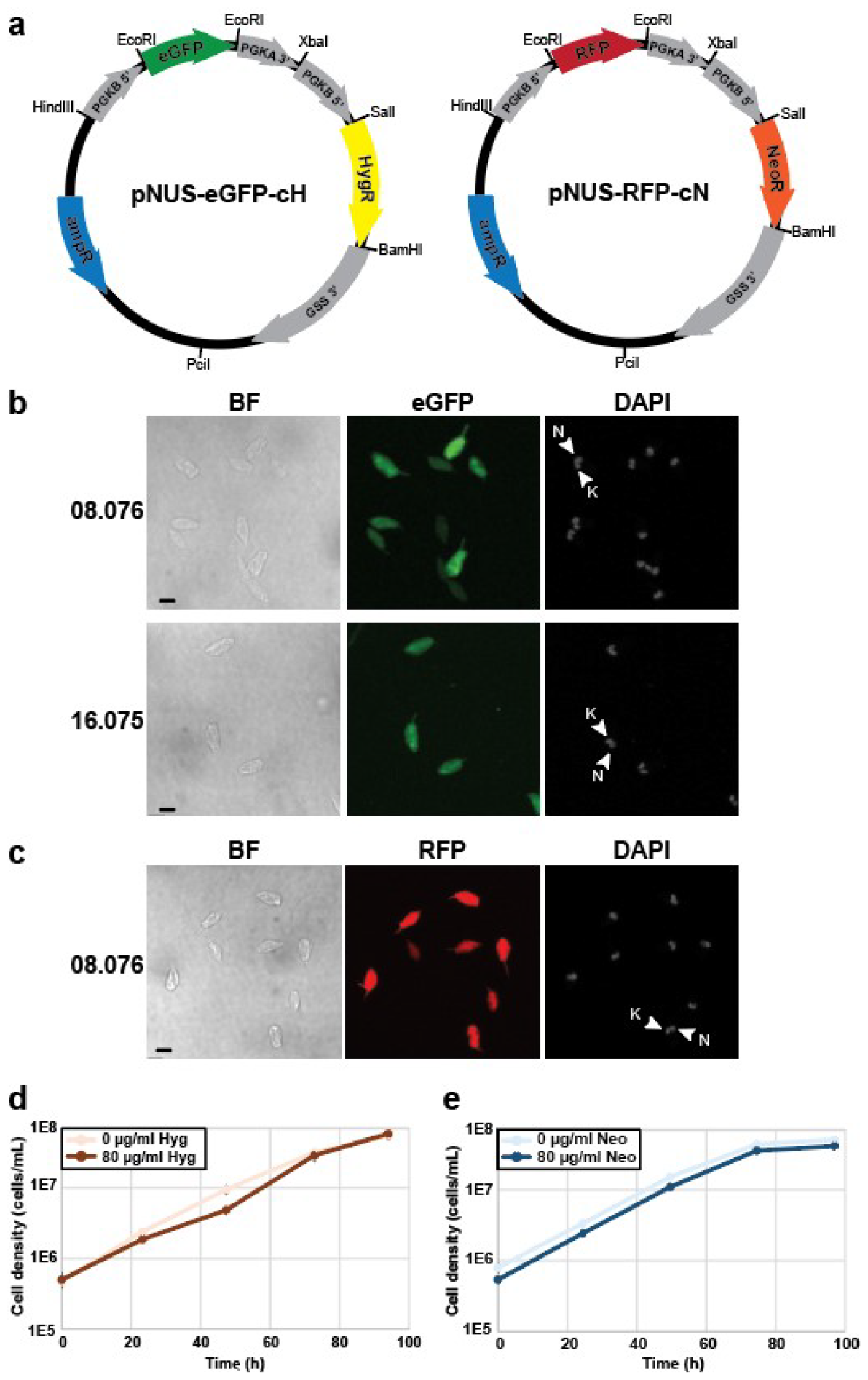
Genetic modification of *C. bombi* cells. **a)** Plasmid map of pNUS-eGFP-cH [24]. The open reading frames for enhanced green fluorescence protein (eGFP), hygromycin resistance (HygR) and ampicillin resistance (ampR, for propagation of plasmids in *E. coli*), as well as 5’ and 3’ UTRs from the phosphoglycerate kinase (PGK) and glutathione synthetase (GSS) genes including sequences for transcript processing are shown. Plasmid map of pNUS-RFP-cN [24] showing open reading frames encoding red fluorescence protein (RFP), neomycin resistance (NeoR), and ampR as well as 5’ and 3’ UTRs. **b)** *C. bombi* strains 08.076 [22] and 16.075 [33] nucleofected with the pNUS-eGFP-cH plasmid. BF, brightfield (visible light); eGFP, enhanced green fluorescent protein signal visible in the green channel; DAPI, DNA stain. Mitochondrial DNA (kinetoplast, K) and the nucleus (N) in a single cell are indicated with arrowheads. Fluorescent images are z-stack maximum projections. Scale bar is 5 μm. **c)** *C. bombi* strain 08.076 harboring the pNUS-RFP-cN plasmid imaged in brightfield (BF), red fluorescence channel (RFP), and with DAPI. K, kinetoplast DNA; N, nucleus. Scale bar is 5 μm. **d)** Growth curve of *C. bombi* strain 08.076 bearing the pNUS-eGFP-cH plasmid in 0 or 80 μg/mL Hyg. For each condition, the mean of three replicates is shown. Error bars are standard error. **e)** Growth curve of *C. bombi* strain 08.076 bearing the pNUS-RFP-cN plasmid in 0 or 80 μg/mL Neo. For each condition, the mean of three replicates is shown. Error bars are standard error.

We used the same procedure to introduce pNUS-GFP-cH into a different *C. bombi* isolate, 16.075 [33] (Fig 2b). As with 08.076, green fluorescence in transformed 16.075 cells was significantly above background (autofluorescence detectable in both isolates imaged with long exposures, Fig S1). Thus, our nucleofection procedure seems generalizable to more than one strain/isolate of culture-adapted *C. bombi*.

One disadvantage of modification by episomal plasmids is heterogeneity of expression, since each parasite can contain a different number of episomes that are imperfectly segregated during cell division [24,27]. Therefore, after several passages, we increased the concentration of Hyg or Neo from 40 μg/mL to 80 μg/mL to select for cells with higher episomal copy number, although expression levels still varied somewhat between cells. To confirm that eGFP-or RFP-expressing cells were fully drug resistant, we performed growth curves and found that, in contrast to parental strains, 08.076 *C. bombi* transformed with pNUS-eGFP-cH or pNUS-RFP-cN showed robust growth in the presence of 80 μg/mL of the appropriate selecting drug (Fig 2d, e).

### (3) Expression of fluorescent proteins with distinct subcellular localizations

*Crithidia bombi*, like other trypanosomatids, are complex eukaryotic cells with a variety of compartments and a distinctly polarized subcellular organization [34,35]. In related species, subcellular organization can change as the parasite undergoes morphological and metabolic adaptation to different environments [35–38]. In addition to changes in cell and organelle shape, the localization of individual proteins can also vary during the cell and life cycle [35,39]. Determining the location of a particular protein can provide important clues to its function. For these reasons, it is useful to have markers for various subcellular compartments to monitor these organelles and to compare to the localization of uncharacterized proteins. For example, the mitochondrion of trypanosomatids is typically an elaborate branched network that extends throughout the cell. The dixenous trypanosomatid *Trypanosoma brucei* dramatically alters both mitochondrial shape and function as it alternates between mammalian and insect hosts [40]. The branched mitochondrial network can resemble other organelles, such as the endoplasmic reticulum, making colocalization with a known marker required to confirm the subcellular location of a protein [35]. Organelle-specific dyes, such as MitoTracker, can be useful for co-localization but they are dependent on membrane potential and dye toxicity, which can complicate imaging of live cells. Mitochondria are metabolic and signaling hubs whose function requires the post-translational import of hundreds of nuclear-encoded proteins. The function and proper localization of many of these proteins are likely required for survival and replication of *C. bombi* parasites.

To label the mitochondrion, we introduced a variation of pNUS-eGFP-cH in which a mitochondrial targeting signal from the related parasite *T. brucei* was fused to the open reading frame (ORF) of eGFP to produce pNUS-mitoeGFP-cH (Fig 3a) [25]. As observed in *C. fasciculata*, introducing this plasmid into *C. bombi* labels the branched tubular mitochondrion, which we verified by co-localization with MitoTracker (Fig 3b).

**Fig. 3.**
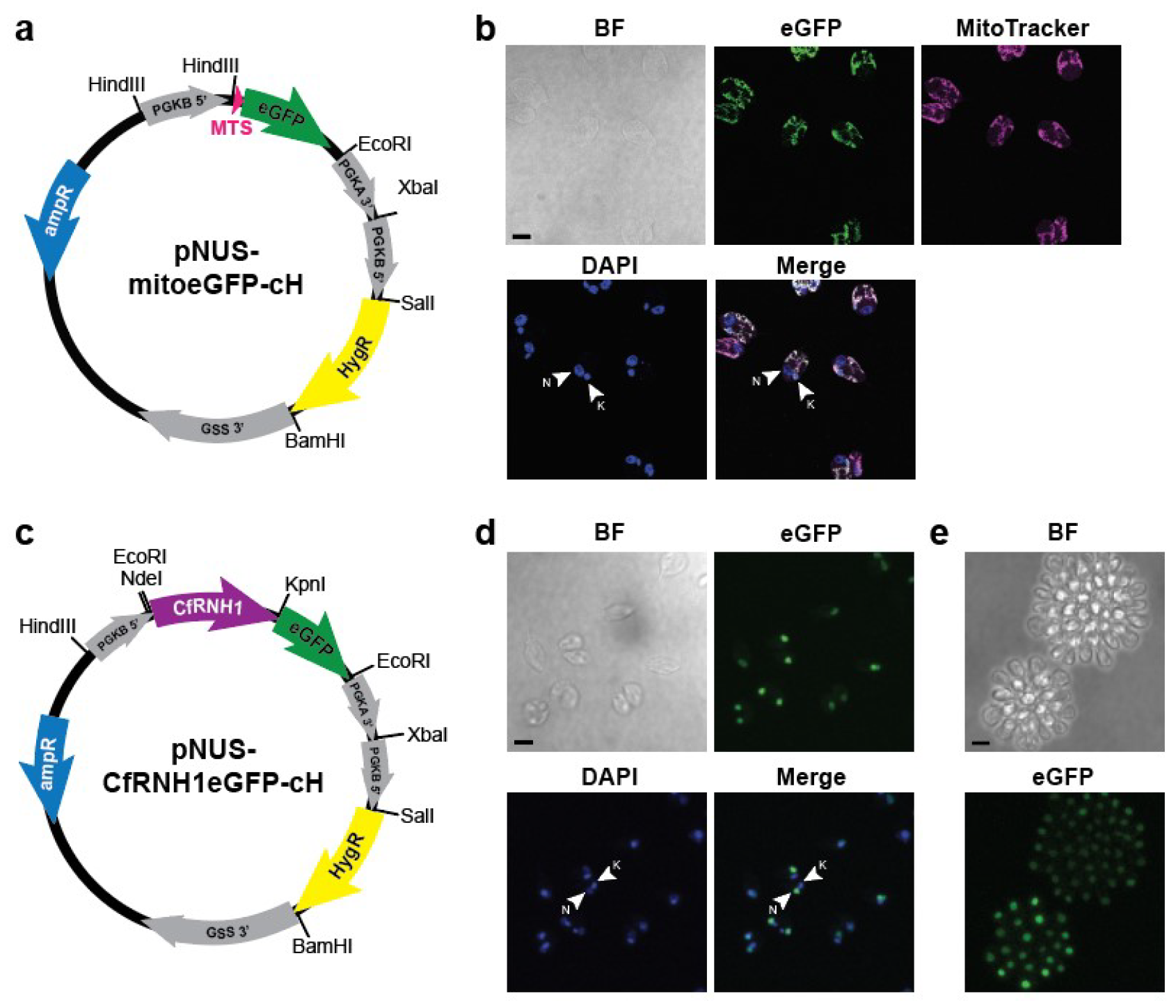
Organelle markers for mitochondria and nuclei in *C. bombi*. **a)** Plasmid map of a construct for episomal expression of a mitochondrion-targeted eGFP [25]. 5’ and 3’ UTRs for transcript processing and stability are shown in gray. PGK, phosphoglycerate kinase; GSS, glutathione synthetase; MTS, mitochondria targeting signal; eGFP, enhanced GFP; HygR, hygromycin resistance; ampR, ampicillin resistance. Some relevant restriction enzyme sites are also shown. **b)** *C. bombi* strain 16.075 [33] nucleofected with pNUS-mitoGFP-cH. BF, brightfield; eGFP, mitochondrial eGFP fluorescence; MitoTracker, membrane potential dependent mitochondrial red fluorescent dye; DAPI, DNA stain (N, nucleus; K, kinetoplast). Merge shows overlay of fluorescent channels. Scale bar is 5 μm. **c)** Plasmid map of pNUS-*Cf*RNH1eGFP-cH. *Cf*RNH1, RNH1 gene from *Crithidia fasciculata*. Other abbreviations as in (a). Some relevant restriction enzyme sites are shown. **d)** Swimming *C. bombi* strain 08.076 [22] nucleofected with the pNUS-*Cf*RNH1eGFP-cH plasmid showing nuclear expression of eGFP-tagged *Cf*RNH1 (eGFP); DAPI, DNA stain (N, nucleus; K, kinetoplast); brightfield (BF); and merge of fluorescent channels. Scale bar is 5 μm. **e)** 08.076 *C. bombi* cells growing as attached rosettes on a MatTek dish and expressing pNUS-*Cf*RNH1eGFP-cH were imaged live in brightfield (BF) and in the green channel to show nuclear localization of *Cf*RNH1eGFP. Scale bar is 5 μm.

While alterations in mitochondrial shape could indicate changes in parasite metabolism, fluorescent nuclei would provide a clear identification of dividing cells. Determining the frequency of dividing cells in different insect tissues and at different stages of the infection would inform models for rates of colonization and infectivity of the insect host. To label the nucleus, we created a plasmid in which the ORF for the RNase H1 gene from *C. fasciculata* (*Cf*RNH1) was fused to eGFP at its C-terminus in pNUS-eGFP-cH to create pNUS-*Cf*RNH1eGFP-cH (Fig 3c). Introduction of this plasmid into *C. bombi* followed by Hyg selection resulted in cell lines expressing GFP in the nucleus as evidenced by co-localization with the DNA stain 4’,6-diamidino-2-phenylindole, (DAPI) (Fig 3d). In *C. fasciculata*, earlier work showed that the gene for *Cf*RNH1 contains alternate start codons allowing for two versions of the protein, one of which contains a mitochondrial targeting signal [41]. In *C. bombi*, the *Cf*RNH1eGFP signal is concentrated in the nucleus, indicating that the dual localization of this enzyme may not occur. However, our construct included only the ORF (including both possible start codons) of *Cf*RNH1 but not the native 5’ processing signal, which may be important for stability of the longer transcript [41].

In order to colonize their insect host, *C. bombi* cells must attach via their flagella to the lining of the hindgut [5]. This allows the parasites to replicate as attached cells without being eliminated by defecation. In *C. fasciculata* and other species, this distinct developmental stage can also occur *in vitro* through contact with tissue culture plastic [23,38,42–46]. To see if this was the case for *C. bombi*, we allowed log-phase parasites to adhere to a plastic dish for 24 hours, followed by washing with 1X PBS to remove non-adherent cells. As in *C. fasciculata*, some cells attached and divided, producing attached groups of cells called rosettes. These rosettes could be imaged by live-cell fluorescent microscopy, allowing for visualization of fluorescent markers in both swimming and attached parasites (Fig 3e).

### (4) *Visualizing parasites in* Bombus impatiens

While genetic modification of *C. bombi* will enable morphological and functional studies *in vitro*, it also has the potential to improve visualization of host-parasite interactions and the progress of infections in the natural host under different conditions. Both 08.076 and 16.075 parasite strains were isolated from *B. impatiens* and were culture adapted for sequence analysis and other studies [22,33]. However, mixed infections containing multiple distinct strains of *C. bombi* are common, and some variability between isolates might be expected [22,47]. In addition, extended passaging in culture could fundamentally change the biology of the organism. Therefore, we sought to modify a clonal isolate of *C. bombi* that had been recently obtained from a laboratory colony of *B. impatiens*. We dissected guts from infected bees, homogenized the tissue, transferred parasites to medium, and obtained clones by limiting dilution in 96-well plates in the presence of antibiotics and antifungals.

We selected one of these clones, WHA1, and performed nucleofection to introduce the pNUS-eGFP-cH and pNUS-RFP-cN plasmids in separate experiments as described above. Following selection of drug-resistant lines, we confirmed fluorescence of cells cultured *in vitro* by microscopy. We then used these parasites (*Cb*-WHA1-GFP and *Cb*-WHA1-RFP) to infect commercial *B. impatiens* (Fig 4) and visualized fluorescent cells *in vivo*. As a negative control, microscopy was also completed on digestive tracts of uninfected *B. impatiens*. While there is some autofluorescence of the bumble bee tissue (Fig S2), particularly in the green channel, fluorescent parasites were clearly seen in the guts of the infected bees, attached as clusters to the lining of the hindgut and rectum.

**Fig. 4.**
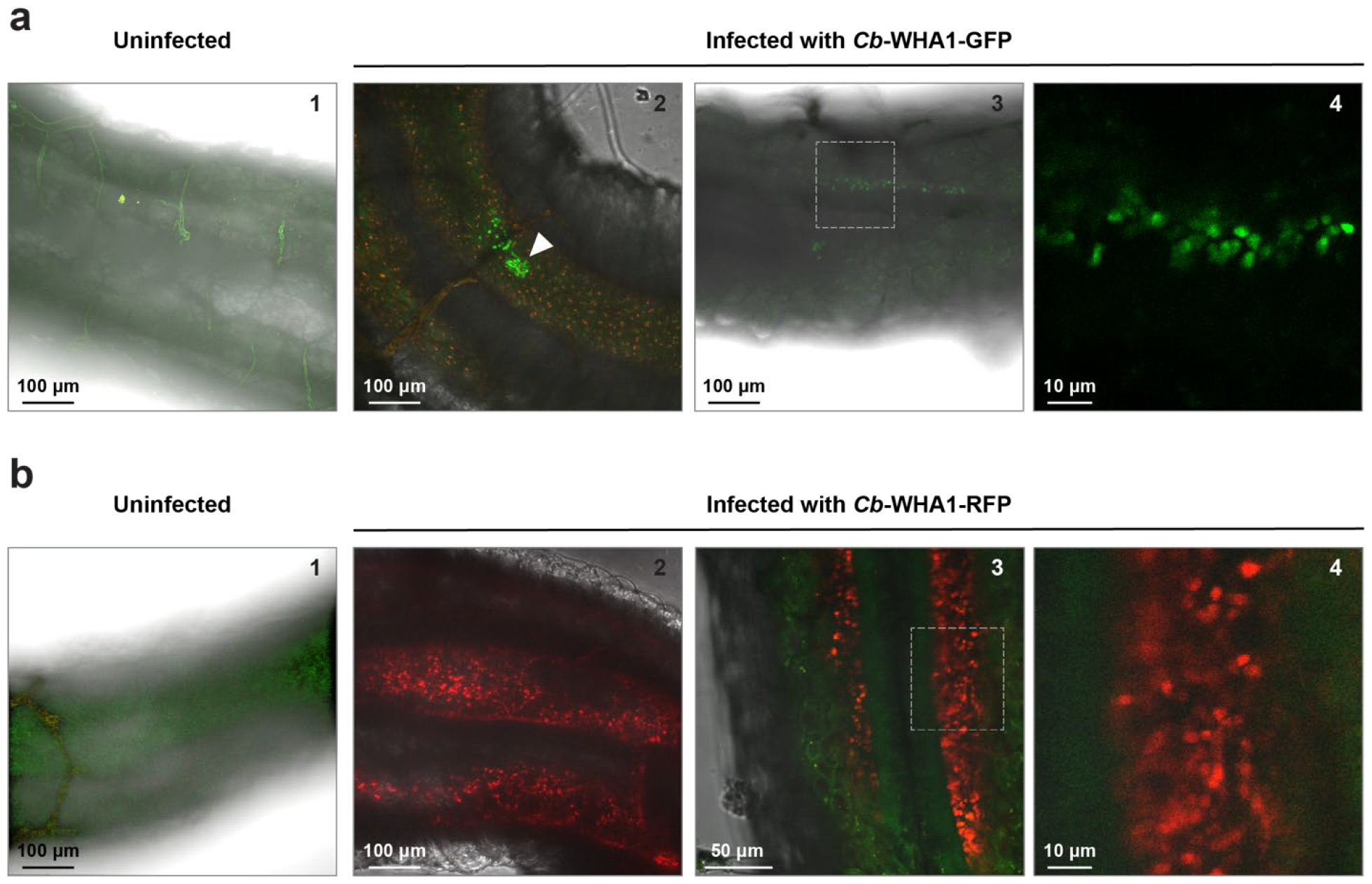
Visualizing fluorescent parasites *in vivo*. **a)** The ileum of bees infected with parasites expressing pNUS-eGFP-cH (panels 2-4) were compared to the ileum of uninfected bees (panel 1). Arrowhead indicates a cluster of eGFP-positive parasites (panel 2). The dotted box in panel 3 shows the area of enlargement in panel 4. Panel 4 shows fluorescence in the green channel only. Panels 1-3 are merges of red and green fluorescence plus brightfield. Scale bars are as shown. **b)** The ileum of an uninfected bee (panel 1) compared to sections of the ileum from bees infected with *C. bombi* parasites expressing pNUS-RFP-cN (panels 2-4). The dotted box in panel 3 shows the area of enlargement in panel 4. All images are z-stack max projections and merges of red fluorescence, green fluorescence, and brightfield. Scale bars are as shown.

## Discussion

We present a method for introducing plasmids into *C. bombi* and selecting modified parasites with two different selectable markers. We have shown that 5’ and 3’ UTRs containing transcript-processing signals derived from *C. fasciculata* function for stable expression in *C. bombi*. In addition, introduction of transgenes bearing localization signals from other trypanosomatids direct similar subcellular localization patterns in *C. bombi*. These modifications improve visualization of cellular morphology of swimming and attached developmental forms, both of which can be generated *in vitro* under standard culture conditions. Genetically modified parasites also retained the ability to infect *B. impatiens*, facilitating detailed studies of host-pathogen interactions.

We successfully modified three independent isolates of *C. bombi*, one of which was isolated directly from a laboratory colony of *B. impatiens* as part of this study. Since the method appears to be both robust and generalizable, it might also be applied to the modification of other pollinator pathogens, such as *Lotmaria passim*. The introduction of plasmids as episomes means that genome sequence data is not required, transfection efficiencies are relatively high, and the same series of plasmids may be used for different species and strains.

We observed that RFP-expressing *C. bombi* displayed much brighter fluorescence than eGFP-expressing cells, allowing for greater sensitivity *in vivo*. This could be due to variation in the amount of correlation between resistance to a particular drug and the average episomal copy number per cell of the selectable marker. Neomycin may select for parasites with larger numbers of episomes per cell, leading to a greater proportion of cells above the threshold for detection. The GFP construct, in contrast, was selected using hygromycin, which may require only a few copies of the resistance gene per cell to allow for growth.

While we have focused on the production of fluorescent parasites to improve sensitivity and resolution for describing infection dynamics *in vivo*, we believe that there are many exciting applications of molecular genetic approaches in these parasites. For example, genome-wide gene expression data obtained through RNA sequencing has revealed transcripts that may enable parasite survival in the host [14]. Labeling these genes with epitope or fluorescent tags would enable researchers to follow their dynamic localization during infection and colonization of the gut. Such tagged proteins could also be used to purify protein complexes that allow parasite adaptation to host microenvironments or that function at the host-parasite interface. In strains for which a genome sequence is available, plasmid constructs could be created to modify genomic loci. This would allow for tagging a gene at its endogenous locus or, compellingly, creating stable genomic knockouts to evaluate gene function in parasite growth and host interactions.

Future research focused on the development of a reliable and quantitative attachment assay for parasites *in vitro* will allow investigators to examine the effects of culture conditions, including floral products, on growth and the developmental switch between swimming and attached forms, a transition that is likely critical for effective colonization of the bee gut. This attachment assay will permit rapid and mechanistic testing of hypotheses that can then be extended to the laboratory infection model and the interpretation of findings in field conditions. We believe that the introduction of molecular genetic tools for manipulation of *C. bombi* will enable integrative approaches across disciplines and scales to begin to bridge the knowledge gaps for how parasites impact bee pollinator health.

## Acknowledgments

This work was supported by a National Science Foundation IntBIO award number 2128223 to LSA and MLP. We thank Ben Sadd and Emmanuel Tetaud for advice and reagents, and Koppert Biological and Biobest for discounted or donated bumble bee colonies. *In vivo* microscopy was performed in the Light Microscopy Facility and Nikon Center of Excellence at the Institute for Applied Life Sciences at the University of Massachusetts Amherst with support from the Massachusetts Life Sciences Center. We especially thank Madeline Malfara along with Lindsay Bair, Laura Anastor-Walters, Gabrielle Schusler, and Nancy Peltier at Villanova University for technical assistance.

## Statements and Declarations

### Funding

This work was supported by a National Science Foundation IntBIO award number 2128223 to LSA and MLP. This work was also supported by a Lotta Crabtree Fellowship from the University of Massachusetts Amherst as well as the CAFE Hatch Award to SKG. This material is based upon work supported by the National Institute of Food and Agriculture, U.S. Department of Agriculture, and the Center for Agriculture, Food and the Environment at University of Massachusetts Amherst, under project number NE2001. The contents are solely the responsibility of the authors and do not necessarily represent the official views of the USDA or NIFA.

### Competing Interests

The authors declare no competing interests.

### Author Contribution

BVB, SGL, SKG, LSA, and MLP conceived of the ideas and designed experiments. BVB, SGL, SKG, FAS, and MLP implemented the experiments and collected the data. BVB, SGL, FAS, SKG, and MLP contributed to visualization. BVB, SGL, SKG, and MLP wrote the first draft of the manuscript. All authors contributed critically to manuscript edits and gave final approval for publication.

### Data Availability

Additional data, including plasmid sequences, are available upon request.

## Notes

### Competing Interest Statement

The authors have declared no competing interest.

