## Supplemental Figures for "Genetic modification of the bee parasite *Crithidia bombi* for improved visualization and protein localization"

### Supplemental Material

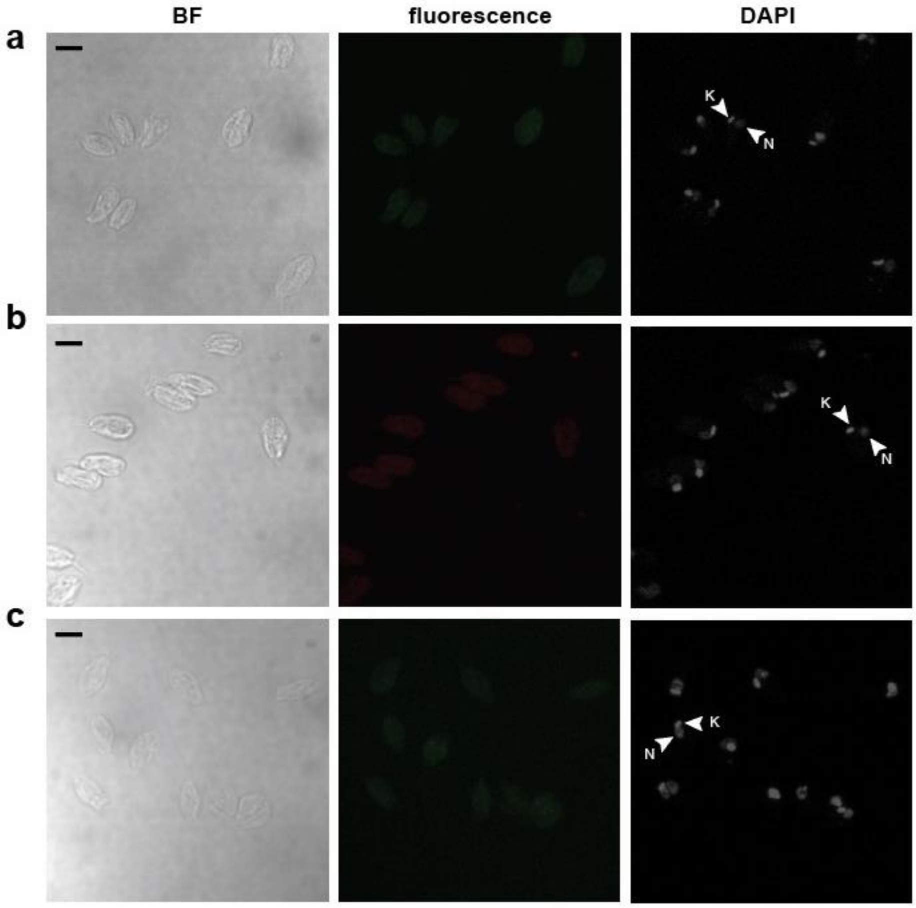


Fig. S1 *Crithidia bombi* parental strains do not show fluorescence. **a)** Parental strain *C. bombi* 08.076 imaged for brightfield (BF), green fluorescence (eGFP) and DAPI. K, kinetoplast; N, nucleus. **b)** Parental strain *C. bombi* 08.076 images for brightfield, red fluorescence, and DAPI. **c)** Parental strain *C. bombi* 16.075 imaged for brightfield, green fluorescence, and DAPI. All images are z-stack maximum projections. Scale bar is 5 µm.

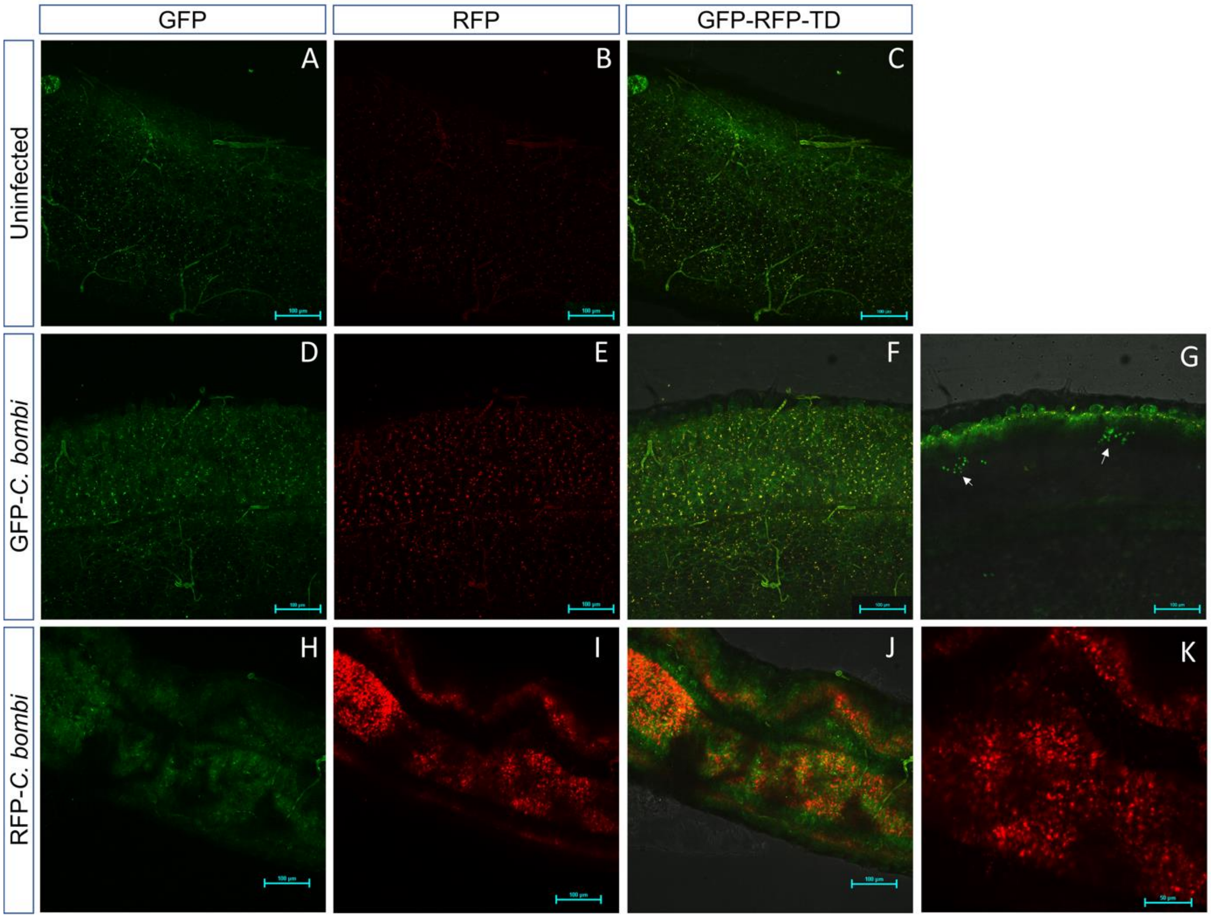


**Fig. S2** *Crithidia bombi* expressing fluorescent transgenes can be imaged *in vivo*. The ileum of uninfected *B. impatiens* **(A-C)** and *B. impatiens* infected with either GFP **(D-G)** or RFP **(H-K)** -expressing *C. bombi* were imaged by confocal microscopy for green fluorescence (GFP), red fluorescence (RFP) and brightfield (TD). In G clusters of parasites are indicated by arrows. Some autofluorescence is detectable in both green and red channels that is distinguishable from fluorescent parasites. Scale bars are as shown.
